# Estimating animal density in three dimensions using capture-frequency data from remote detectors

**DOI:** 10.1101/852475

**Authors:** Juan S. Vargas Soto, Rowshyra A. Castañeda, Nicholas E. Mandrak, Péter K. Molnár

## Abstract

1. Remote detectors are being used increasingly often to study aquatic and aerial species, for which movement is significantly different from terrestrial species. While terrestrial camera-trapping studies have shown that capture frequency, along with the species’ movement speed and detector specifications can be used to estimate absolute densities, the approach has not yet been adapted to cases where movement occurs in three dimensions. Frameworks based on animal movement patterns allow estimating population density from camera-trapping data when animals are not individually distinguishable.
2. Here we adapt one such framework to three-dimensional movement to characterize the relationship between population density, animal speed, characteristics of a remote sensor’s detection zone, and detection frequency. The derivation involves defining the detection zone mathematically and calculating the mean area of the profile it presents to approaching individuals.
3. We developed two variants of the model – one assuming random movement of all individuals, and one allowing for different probabilities for each approach direction (e.g. that animals more often swim/fly horizontally than vertically). We used computer simulations to evaluate model performance for a wide range of animal and detector densities. Simulations show that in ideal conditions the method approximates true density well, and that estimates become increasingly accurate using more detectors, or sampling for longer. Moreover, the method is robust to invalidation of assumptions, accuracy is decreased only in extreme cases where all detectors are facing the same way.
4. We provide equations for estimating population density from detection frequency and outline how to estimate the necessary parameters. We discuss how environmental variables and species-specific characteristics affect parameter estimates and how to account for these differences in density estimations.
5. Our method can be applied to common remote detection methods (cameras and acoustic detectors), which are currently being used to study a diversity of species and environments. Therefore, our work may significantly expand the number and diversity of species for which density can be estimated.

## Introduction

Remote detection methods like camera-traps originated for studying surface-dwelling species but are increasingly being used to study aquatic, arboreal, and airborne species. These technologies require considerably less effort and resources than alternatives like transects, particularly for long-term studies, and are less invasive. In aquatic studies, remote underwater video has allowed more efficient, extensive, and less biased sampling than diver-based surveys (King, George, Buckle, Novak, & Fulton, 2018; Mallet & Pelletier, 2014). Similarly, acoustic detectors are becoming a more accessible and popular tool for studying and monitoring a variety of taxa such as birds (Blumstein et al., 2011; Celis-Murillo, Deppe, & Allen, 2009), echolocating bats (Marques et al., 2013), and even fishes that produce species-specific sounds, especially with the aid of automatic identification algorithms (Lindseth, Lobel, Lindseth, & Lobel, 2018). Camera-traps in turn have been useful to study the demographics of elusive species or to learn about biodiversity across different spatial scales (Ahumada et al., 2011; Barea-Azcón, Virgós, Ballesteros-Duperón, Moleón, & Chirosa, 2007; Karanth, Nichols, Kumar, & Hines, 2006). In every case, the technologies, software and analysis methods are constantly evolving (Burton 2015).

Two decades of theoretical developments have created a vast literature of methods for estimating the density of wild populations from camera-trapping data, and some of these methods can be adapted to other types of detectors. Mark-recapture methods, for example, have been used to study aquatic species with individually recognizable spots like whale sharks (*Rhincodon typus*) and eagle rays (Myliobatidae) (González-Ramos, Santos-Moreno, Rosas-Alquicira, & Fuentes-Mascorro, 2017; Meekan et al., 2006). However, most species lack individual markings and are therefore not amenable to these approaches (e.g., Karanth et al. 2006). For these species, existing frameworks for estimating density either provide relative estimates only [e.g. detection frequency, minimum number of detected individuals (Sherman, Chin, Heupel, & Simpfendorfer, 2018), maximum number of conspecifics in a single frame (Willis, Millar, & Babcock, 2000)], or make assumptions about how animals move for estimating absolute density [e.g. movement around a home range centre (Campos-Candela, Palmer, Balle, & Alós, 2018), random walks (Nakashima, Fukasawa, & Samejima, 2018), ideal gas movement (Rowcliffe, Field, Turvey, & Carbone, 2008)]. These movement-based frameworks need to be adapted for key differences between terrestrial and aquatic or aerial species that perceive the world and move in three dimensions.

One approach for estimating population density when individuals cannot be distinguished is the Random Encounter Model (REM), formalized for encounters between animals and camera-traps by Rowcliffe et al. (2008). The model assumes that animals move like ideal gases (in straight lines, in random directions, with constant speed) and as a result the frequency at which a species is photographed by camera-traps (henceforth, “detection frequency”) scales positively with the number of individuals present in an area (i.e. population density), the species’ mean speed, and the size of the camera’s detection zone. This relationship can therefore be used to estimate density from the detection frequency. While this method requires more information about a study species than relative abundance measures, it considerably expands the number of species for which density can be estimated using camera-traps. The REM framework can be adapted to species that move in three dimensions rather than two, considering the three-dimensional shape and size of the detection zone.

Here, we present such an adaptation to estimate absolute density for aquatic species, using underwater cameras, and for birds and echolocating species, using acoustic sensors. Our adaptation substantially expands possibilities for estimating density from remote detection methods when species can be identified but individuals cannot. This method only requires the detection frequency of a species, information about the sensor’s detection zone, and an estimate of the species’ speed. We provide two alternatives; the first assumes completely random movement (within the specifications of the ideal gas framework, cf. below), while the second allows accounting for biases in movement direction (e.g. that fish more often swim parallel, rather than perpendicular, to the bottom). First, we will explain the framework in detail, highlighting the importance of an accurate mathematical description of the detection zone. We will then develop the formulae for density for two types of detectors and test our estimator’s performance using computer simulations.

## Materials and Methods

### 1. Model Derivation

#### 1.1. The Random Encounter Model framework

The REM method for estimating density of unmarked animals from camera-trapping data is based on the idea that the encounter rate between a stationary detector and animals of a given species scales linearly with the species’ density (see Hutchinson & Waser, 2007), with a scaling factor that depends on the species’ movement characteristics and the detector’s ability to record animals that are passing by at various distances (Rowcliffe et al., 2008). This relationship is summarized as:

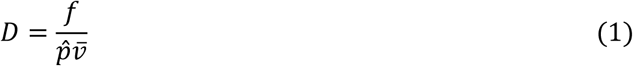

where *D* is animal density, *f* is detection frequency, and *v̄* is mean animal movement speed (See Table 1 for description of parameters). The scaling constant *p̂* can be thought of as the mean distance (in 2D) or area (in 3D) sampled instantaneously by the detector over all possible directions, and therefore depends on the sensor’s technical specifications (opening angles and maximum detection distance) and on environmental variables (e.g. water clarity, foliage or clutter). Here, we will derive an equation for *p̂* for the three-dimensional case. The main steps are 1) determining the shape of the detection zone, 2) describing it mathematically, and 3) calculating the mean area of its two-dimensional projection. The result is used directly in equation 1 to estimate density from capture frequency. We explain in more detail the projection of the detection zone in Section 2 and derive the density estimator in Section 3. We then test the performance of our estimator with computer-simulated capture data.

**Table 1.** Parameters used in the derivation of the 3D random encounter method for estimating animal densities.

| Parameter | Description |
| --- | --- |
| $\hat{p}$ | Mean area of the 2D projection of the 3D detection zone. |
| $S, S_i$ | Surface area of the detection zone, and of its components $i$. |
| $s$ | Maximum distance from the detector at which animals can be detected |
| $\phi$ | Opening angle of the detector, corresponding to the diagonal field of vision in a camera. |
| $\kappa, \lambda, \gamma$ | Horizontal, vertical and diagonal fields of vision of a cropped (camera) detector. |
| $\mu, \nu$ | Opening angles of the disc sectors that make up the sides of a camera's detection zone |
| $\omega, \theta$ | Angles describing the direction of an animal approaching the detection zone, relative to the detector's direction. |
| $D$ | Population density [animals per unit volume] |
| $\bar{v}$ | Mean movement speed of the study species |
| $f$ | Detection frequency: the number of independent recordings of a species by a detector, divided by deployment time |

#### 1.2. Profile of the detection zone, extension from 2D to 3D

For terrestrial camera-traps, where animals move in two dimensions only, *p̂* in equation 1 is the mean profile width of the camera’s detection zone, as presented to approaching animals. Given the shape of the detection zone, the profile width depends on the direction of approach (Fig. 1). While not explicitly stated by Rowcliffe *et al*. (2008), the “profile” is the projection of the detection zone onto a line perpendicular to the direction of approach. One can therefore use Cauchy’s surface area theorem (Cauchy, 1841) to calculate the mean profile width. This theorem states that the average projected area of a convex body is proportional to its surface area. The mean profile width of the two-dimensional detection zone is its perimeter *P* multiplied by a constant:

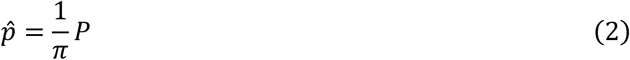

which is the same as equation 2 in Rowcliffe et al. (2008). The theorem can readily be applied to determine the mean profile of a three-dimensional detection zone.

**Figure 1.**
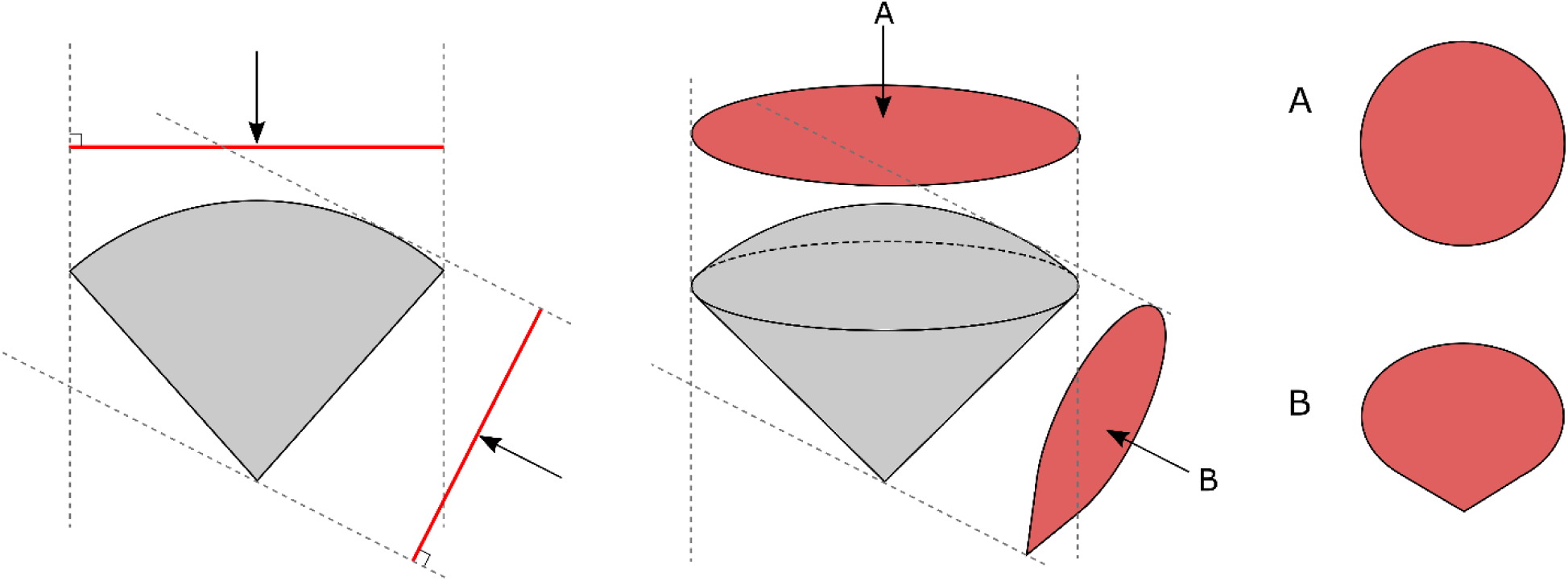
The different profiles presented by the detection zone of a remote detector to an approaching animal in two (left) or three dimensions (center). In 2D, profiles are lines, while in 3D they are surfaces. The silhouettes A and B (right) show the profiles presented to individuals approaching from directions A and B, respectively.

In the case of a three-dimensional detection zone, equation (1) still holds, with the difference that *p̂* corresponds to an area instead of a length. This area is nonetheless the projection of the detection zone onto a lower-dimensional object, i.e. onto a plane. According to Cauchy’s theorem, the mean projected area of a three-dimensional object is obtained from the surface area *S* as:

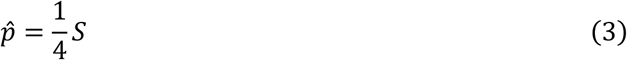

(Cauchy, 1841; Vouk, 1948). To calculate S, we will define the detection zone for two types of sensors: a generic sensor with a conical detection zone (acoustic detectors), and the special case of a sensor with a cropped image (i.e. cameras). For acoustic detectors we also determined how to calculate density when there is bias in directions of approach relative to the direction of the detector, for example bats flying most frequently horizontally into a detection zone facing up.

### 2. Definition of the detection zone

A remote detector’s detection zone is generally determined by two parameters: detection angle and maximum distance from which a signal is detectable. Geometrically, this translates into a cone with a convex base. Acoustic detectors report signals no matter where in this zone they occur, but for cameras some near-boundary signals are lost when images are cropped to a rectangular frame. As such, acoustic detectors have the full “conical-with-hat-shaped” detection zone (Fig. 2A), whereas cameras have a subset of this region defined by the horizontal and vertical angles of view (Fig. 2D). In Section 3.1, we derive expressions to calculate the surface area for both detector types and use Cauchy’s theorem to calculate the respective mean profile area. This method applies if we assume that all directions of approach are equally likely. In section 3.2, we relax this assumption and consider potential biases in the direction of approach.

**Figure 2.**
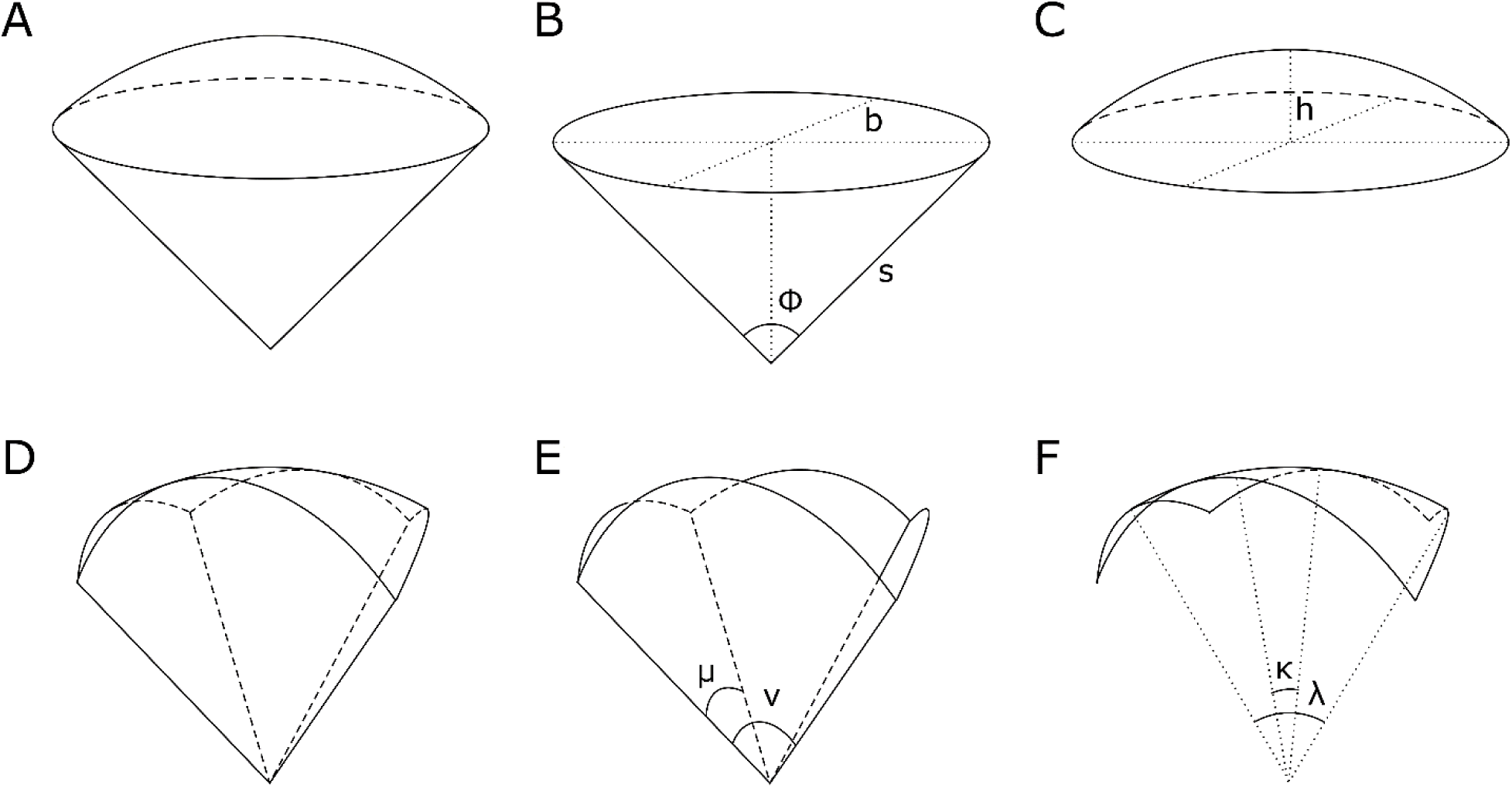
Perspective of the detection zone of (A) an acoustic detector and (D) a camera, where the image is cropped into a rectangle. We decomposed both types of detection zone into bottom and top sections to calculate their surface areas: for acoustic detectors, we considered the lateral surface area *S*_*S*_ of a cone (B), and the area *S*_*C*_ of a spherical cap (C); for cameras, we considered the surface area *S*_*L*_ of four disc segments (E) and the area *S*_*R*_ of a spherical rectangle (F). Parameters are *φ*: opening angle of the detector, *s*: slant height of cone, *b*: basal radius of the cone, *h*: height of the spherical cap, *μ* and *ν*: angles of the disc segments that make up the sides of the cropped detection zone, *κ* and *λ*: horizontal and vertical fields of vision of the camera

#### 2.1. Uniformly distributed approach directions

To calculate the surface area of an acoustic sensor’s detection zone, we divide the detection zone into two components: a conic base (Fig. 2B) and a spherical cap (Fig. 2C). The lateral surface area of the cone is calculated from the slant height, *s* (i.e. the maximum detection distance), and the cone’s basal radius, *b*, as:

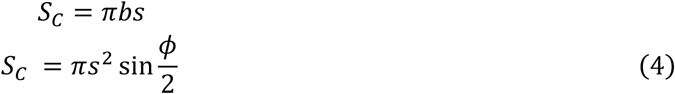

where *φ* is the detector’s opening angle. The spherical cap’s surface area is *S*_*S*_= 2*πsh* (Kern & Bland, 1938), where *h* is the cap’s height, given by 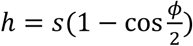 (Appendix A1), so:

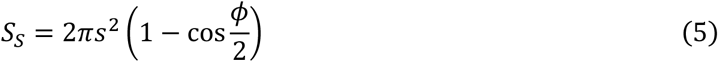

Combining equations (4) – (5), we obtain the mean profile area of an acoustic detector’s detection zone:

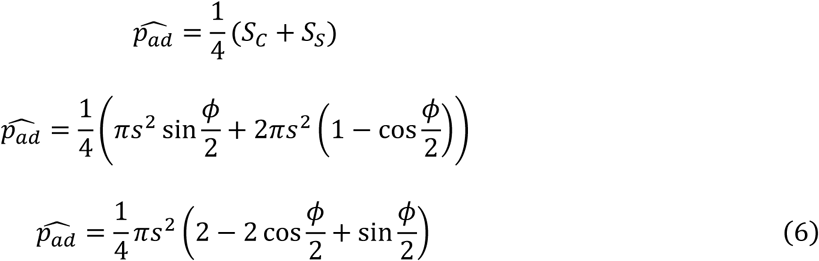

Cauchy’s theorem is only valid for convex bodies, so equation 6 only applies when the detection angle is smaller than *π* (180°) or equal to 2*π* (360°). The latter occurs when signals can be detected from any direction, as is the case with omnidirectional microphones. Consider for example an acoustic detector with a detection angle of 90° (π/2 rad) that can detect signals from 10 meters away. Substituting *s* and *ϕ* in eq. 6 we obtain:

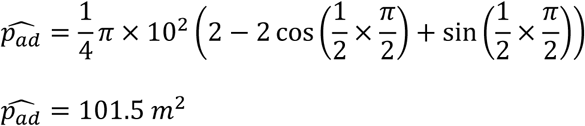

To calculate the surface area of a camera’s detection zone, we note that image cropping creates a detection zone bounded by a spherical rectangle cap sitting atop two pairs of disc sectors of radius *s* and delimited by angles *μ* and *ν* (Fig. 2D). These angles are related to the vertical (*λ*), horizontal (*κ*), and diagonal (*γ*) fields of vision of the camera, of which at least one is normally provided by the manufacturer (see Appendix A3 for how to calculate the unknown angles). As the surface areas of the disc sectors are given by 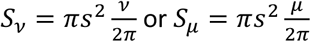, respectively, the total lateral surface area of the camera’s detection zone, S_*L*_ (Fig. 1E), becomes

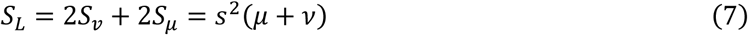

Furthermore, the surface area of the spherical rectangle cap, *S_R_* (Fig. 1F), is obtained by multiplying the surface area of a sphere of radius *s* by the proportion of the sphere occupied by the spherical rectangle:

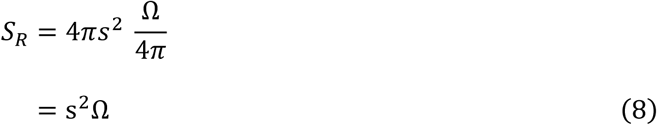

where Ω is the solid angle delimited by the horizontal and vertical fields of vision, *κ* and *λ*, given by (Appendix A2):

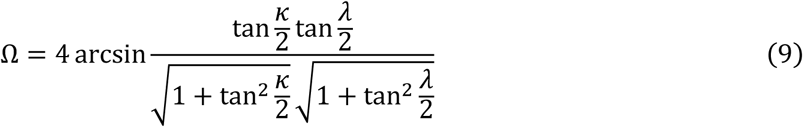

The mean profile area for a camera is therefore:

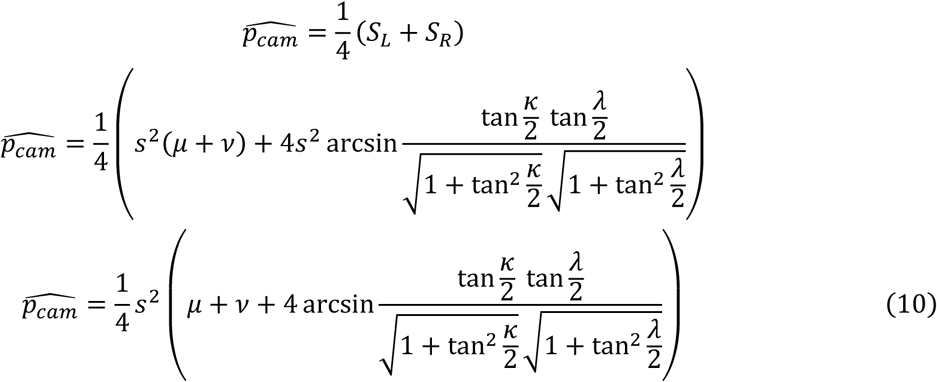

Consider for example an underwater camera with horizontal and vertical fields of vision (FOV) of 122.6° (2.1 rad) and 94.4° (1.6 rad), respectively (these correspond to a GoPro Hero 7 using a wide 4:3 aspect ratio). Using eq. S3.2 we obtain a diagonal FOV of 154.5° (2.7 rad) and using eq. S3.4 we obtain the lateral angles *μ* = 142.1° = 2.5 rad and *ν* = 125.2° = 2.2 rad. Substituting these values in eq. 10, assuming again a detection distance of 10 m, we obtain the mean profile area for this camera:

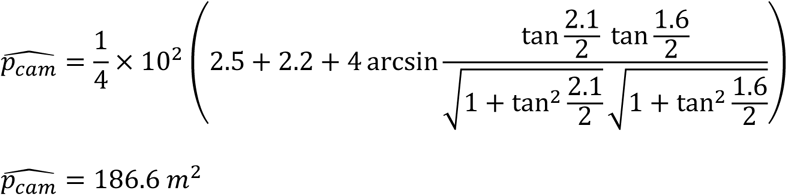

#### 2.2. Bias in direction of movement

The method described above for calculating the detection zone’s mean profile area *p̂* assumes that every direction of approach is equally likely. However, some angles of approach could occur more frequently than others depending both on the study species and the placement and orientation of the detector. Picture, for example, fish moving in a stream in the direction of the current. The profile presented to all of them by a camera is the same and the effective mean profile area is greater or lower than the expected mean profile area, depending on which way the detector is facing. To account for these biases, we derive formulae to calculate the detection zone’s projected area *p* for any direction of approach and then weight these according to the probability of approach directions. As these calculations quickly become lengthy and complicated, we here summarize the approach for acoustic detectors and provide detailed calculations in Appendix A4.

The mean profile area is given by:

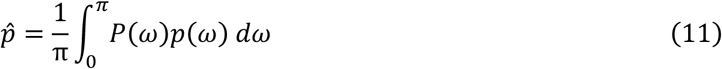

where *P(ω)* is the probability that an individual approaches the detector at angle *ω*, and *p(ω)* gives the profile area corresponding to that direction. Depending on *ω* – the angle relative to the direction of the detector – the different components of the detection zone may be visible or hidden. We obtain therefore four different formulae to calculate *p(ω)*:

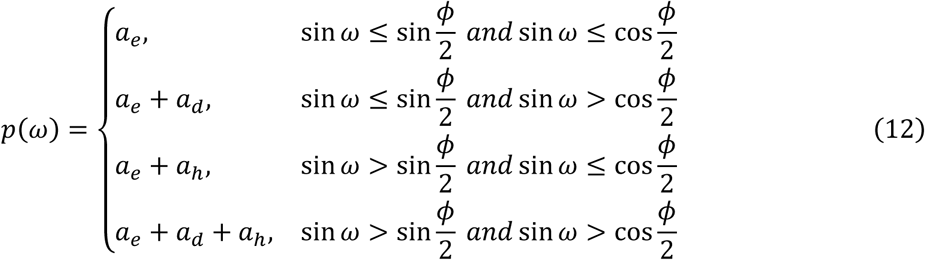

The first element, *a*_*e*_, corresponds to the area of the projection encompassed by the base of the cone (area I in Fig. 3), while the areas *a*_*h*_ and *a*_*d*_ are the projections of the visible parts of the spherical cap (area II) and the cone (area III). The derivation of *a_e_*, *a_h_*, *a_d_*, and the rationale behind them are given in Appendix A4.

**Figure 3.**
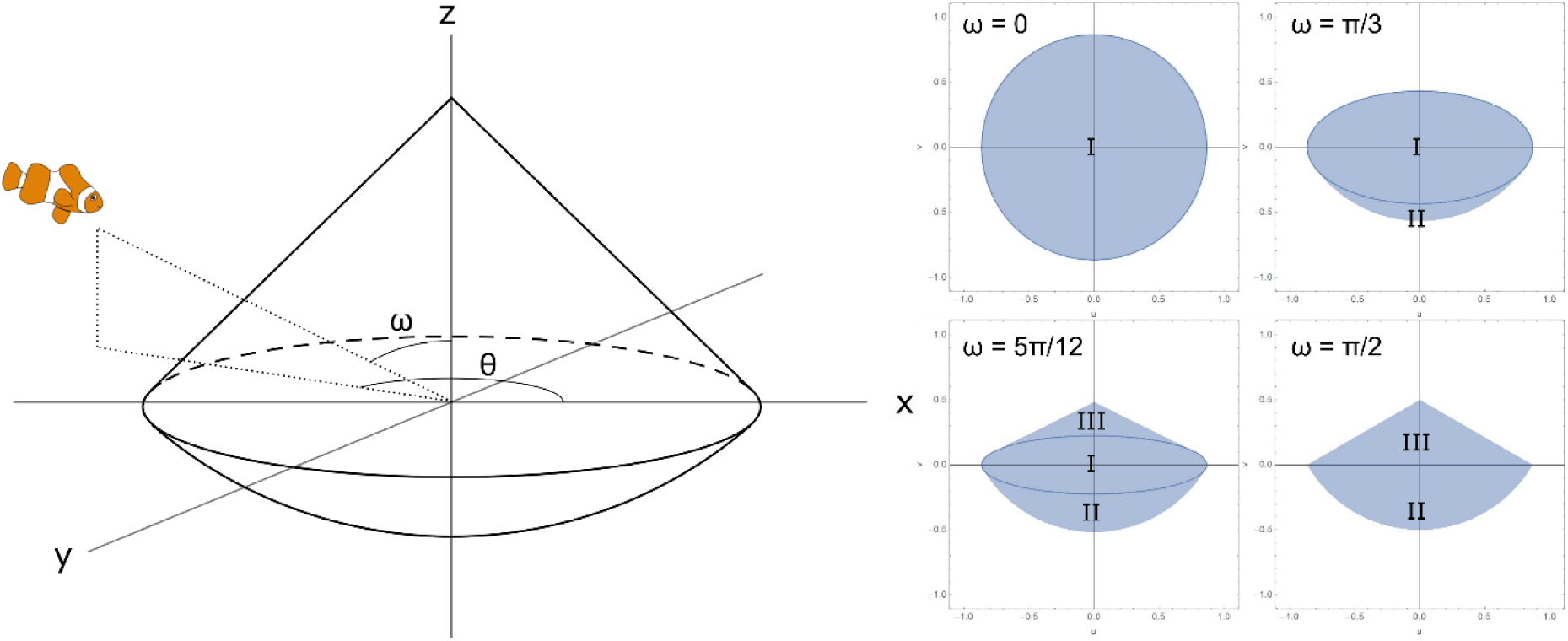
Perspective of a conic detection zone (left), showing the angles *ω* and *θ* that define the direction of approach of an individual. Different angles *ω* result in different profiles (right). The labels indicate the different areas that need to be calculated in each case; I is the area of the projection encompassed by the circle at the base of the cone, II is the projection of the spherical cap, III is the projection of the cone’s sides.

### 3. Simulation tests

We tested the formula for estimating density from detection frequency using computer simulations. Firstly, these serve to confirm that the method performs well under ideal conditions (i.e. when all model assumptions are met and perfect information about the species’ movement is available) and secondly to test the robustness to invalidation of assumptions. The assumptions regarding animal movement are those of the ideal gas model: individuals moving in a straight line, in random directions, and at a constant speed. We assumed perfect detection and an exact knowledge of the detection distance and opening angle and evaluated the method’s performance for a range of animal densities and detector numbers. Furthermore, to determine the robustness of our method, we evaluated its performance for the following scenarios: (i) allowing variation in speed by randomly selecting different individual speeds; (ii) allowing a non-random distribution of approach directions by randomly selecting individual ‘tilt’ angles, combined with realistic scenarios of detector placement and orientation.

We set up the simulation as follows: Individuals were distributed at random locations within a cube of side 10, and each one was assigned a random direction (x, y, z vector components drawn from a uniform distribution from -1 to 1). All individuals moved at the same speed and bounced back into the cube if they reached the reflective boundaries. We tested the density estimator for a range of densities between 0.1 and 10 ind.uv^−1^ (individuals per unit volume), i.e. between 100 and 10000 individuals.

We placed between 5 and 25 detectors facing in random directions at random locations within the ‘sampling zone’, a cube of side 4 situated at the centre of the larger cube. We set a detection distance of 0.5 ud (unit distance) and a detection angle of 45°, which yields a mean profile area of 0.105 ua (unit area) (equation 6).

We set movement speed equal to one length of the detection radius per time step and ran each simulation for 40 steps. We counted an encounter whenever an individual entered a detection zone. Thus, we obtained for each step and each detector the total number of detections up to that time point. We calculated the detection frequency *f* by dividing this cumulative count by the number of steps and then divided *f* by the mean profile area and the movement speed to estimate density (eq. 1). We recorded the mean estimated density across all detectors at every time step. We also determined how performance changed with effort both in terms of the number of detectors and sampling time.

To test how variability in speed among individuals affected density estimates, we ran simulations assigning each individual a speed drawn at random from a normal distribution with mean 0.5 ud/step, and with standard deviation between 0 and 0.1 ud/step. Similarly, to determine the effect of having biased movement directions, we drew the vertical (z) component of the individual direction vectors from a truncated normal distribution centred around 0 and bounded between -1 and 1, so that most individuals would move near horizontally. We assessed the effect of this bias on the performance of the estimator for several scenarios of detector distribution (regular spacing of detectors in 2D on a plane, regular spacing in 3D throughout the sampling cube, random distribution in 3D) and orientation (all horizontal, all vertical, random). These simulations were run with ten detectors.

We iterated each scenario 100 times, and assessed model performance by calculating the bias, precision and accuracy of the density estimate relative to the real density at the end of each simulation. We used the scaled mean error (SME), the coefficient of variation (CV) and the scaled mean square error (SMSE) as indicators of bias, precision, and accuracy, respectively:

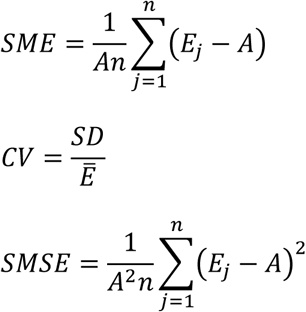

where *Ē* is the mean density estimate, *E_j_* is the density estimate of the *j*^th^ iteration, SD is the standard deviation of density estimates across iterations, *A* is the real density, and *n* is the number of iterations (Walther & Moore, 2005). The bias metric SME indicates whether the true value is over- or underestimated, the precision metric CV indicates the variability among estimates, and the accuracy metric SMSE measures both bias and precision in a single index. The closer to zero each index is, the better the performance of the estimator.

## Results

Simulations show that, under ideal conditions, the estimated density closely approximates the real density. Regardless of the number of detectors used, the estimate density was within 5% of the real value at the end of each simulation (Fig. 4A), and the standard deviation was no larger than 30% of the mean estimate (Fig. 4B). The overall performance of the method nonetheless depended on sampling effort, both in terms of the number of detectors and the sampling time. The lowest number of detectors yielded the greatest bias and the lowest precision and accuracy, and all indices improved substantially with the deployment of additional detectors (Fig. 4A-C). Moreover, bias also decreases, and precision and accuracy increase, as sampling time increases (Fig. 4D-F), meaning that in real-life applications a low availability of detectors could at least be partially compensated for by longer sampling times.

**Figure 4.**
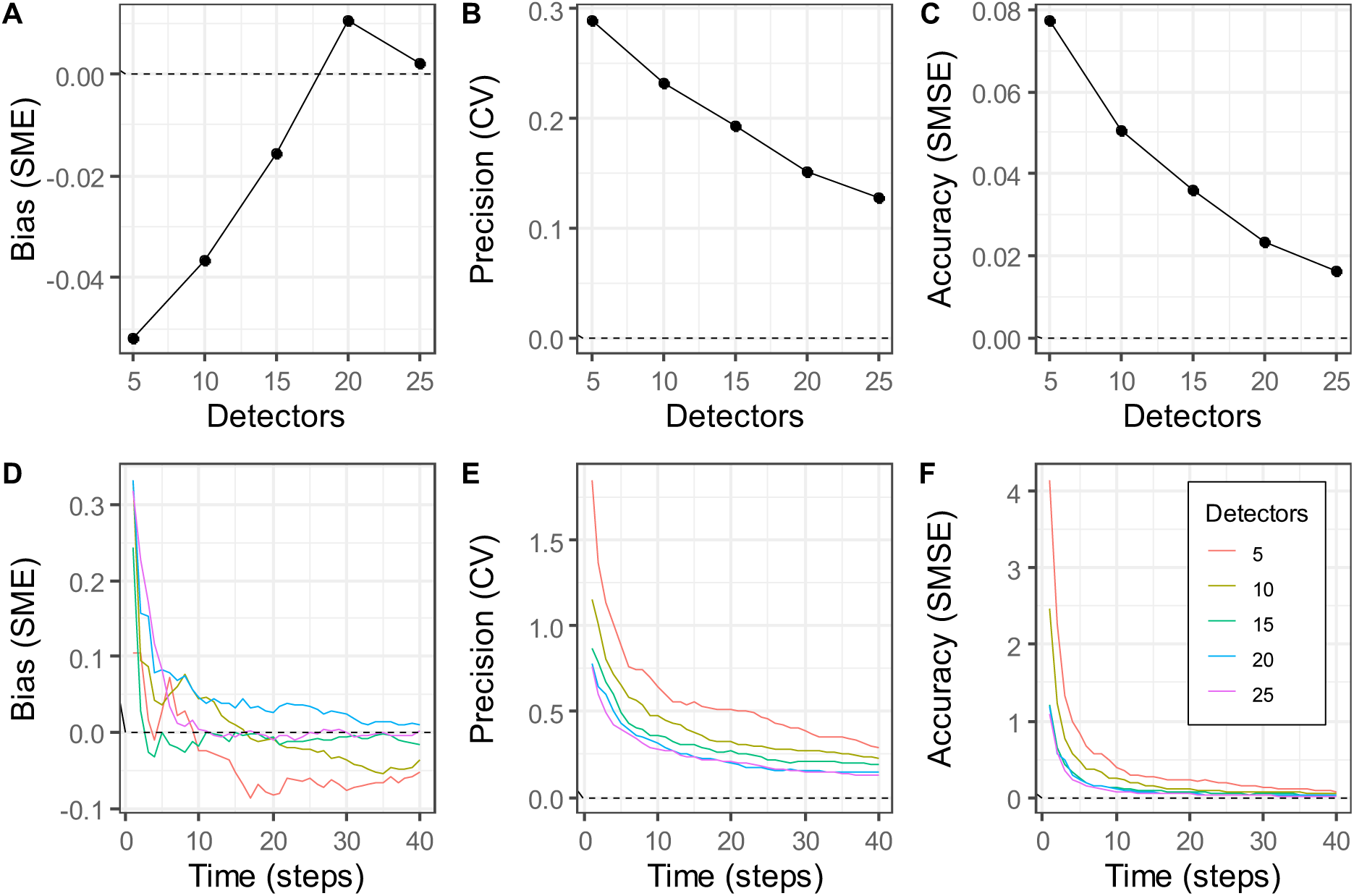
Performance of the 3D density estimation method for different levels of effort. The top row shows the mean bias, precision, and accuracy metric values at the end of the simulations as a function of the number of detectors deployed. The bottom row shows the change in these metrics as time progresses in the simulation.

The real density did not seem to bias the estimator, except at extremely low densities (Fig. 5A). Precision (Fig. 5B), and thus overall accuracy (Fig. 5C), however, depended strongly on the population’s density, with the estimator’s CV decreasing by a factor of more than 9 between the lowest (0.1 ind.uv^−1^) and highest (10 ind.uv^−1^) densities considered.

**Figure 5.**
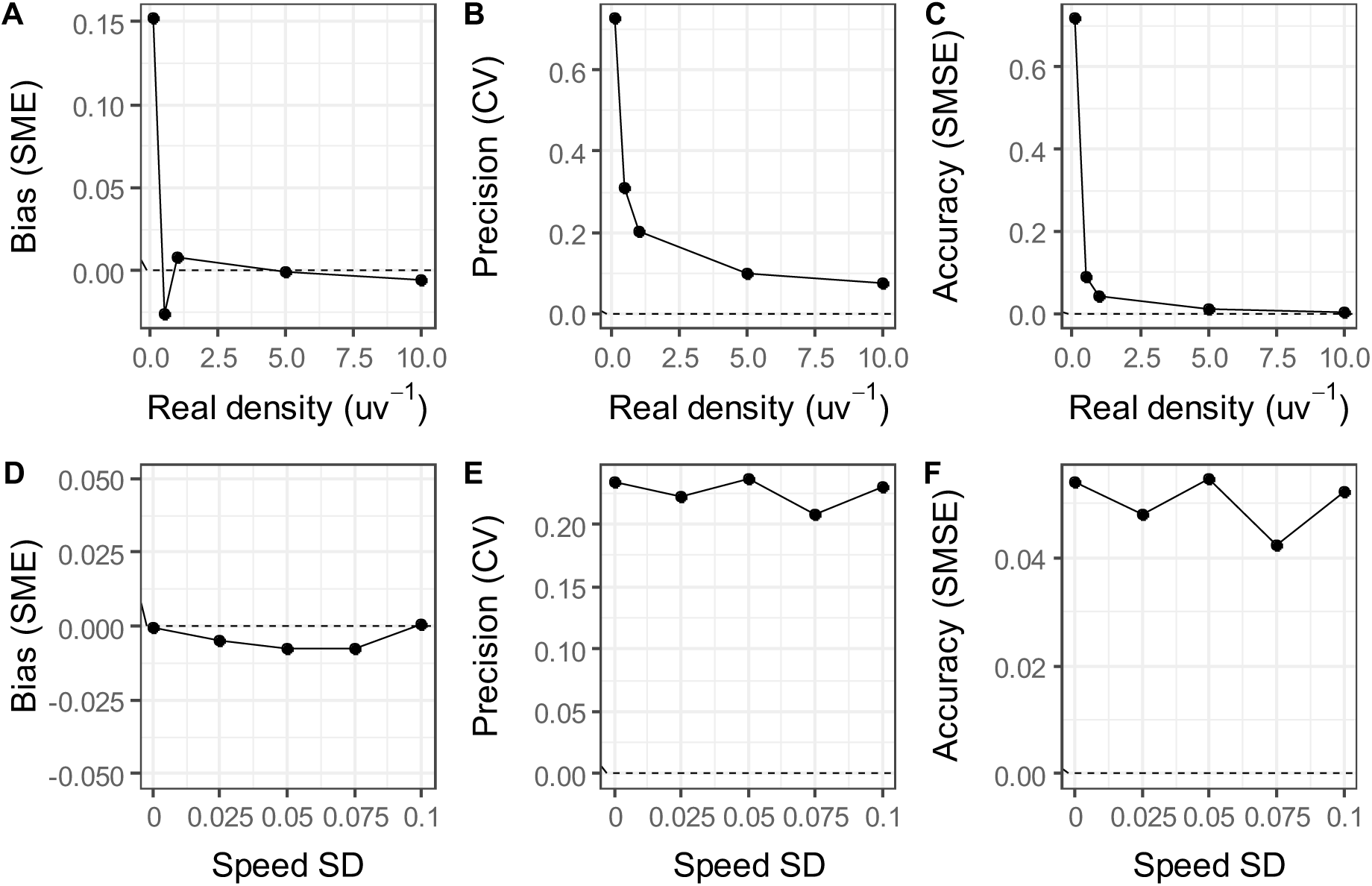
Effects of population density (top row), and among-individual variability in movement speed (bottom row) on the performance of the 3D density estimation method. All parameters were as described in the text, using in particular a constant mean speed of 0.5 ud/step in the top row, and varying the speed among individuals by drawing from a normal distribution with standard deviation between 0 and 0.1 ud in the bottom row.

The simplifying assumption of equal movement speeds among individuals appears to be robust, as introducing among-individual variance in speed did not noticeably affect the estimator’s bias, precision, or accuracy (Figs. 5D-F).

Having biases in the direction of movement, however, did decrease performance in some scenarios, particularly when all individuals were moving horizontally, i.e. with a standard deviation of zero around the mean direction. In this case, precision and accuracy decreased when detectors were placed on a single plane and oriented vertically (Fig. 6D, G) or horizontally (Fig. 6E, H). Conversely, detectors distributed regularly or randomly across the 3D space yielded similar accuracy and precision estimates for all cases of movement direction bias. Nevertheless, there was virtually equal performance regardless of detector placement when detectors faced in random directions (Fig. 6F, I). Moreover, the random orientation of detectors also generally lowered the estimator’s bias compared to the scenarios where all detectors were facing vertically or horizontally (contrast Figs. 6A-C).

**Figure 6.**
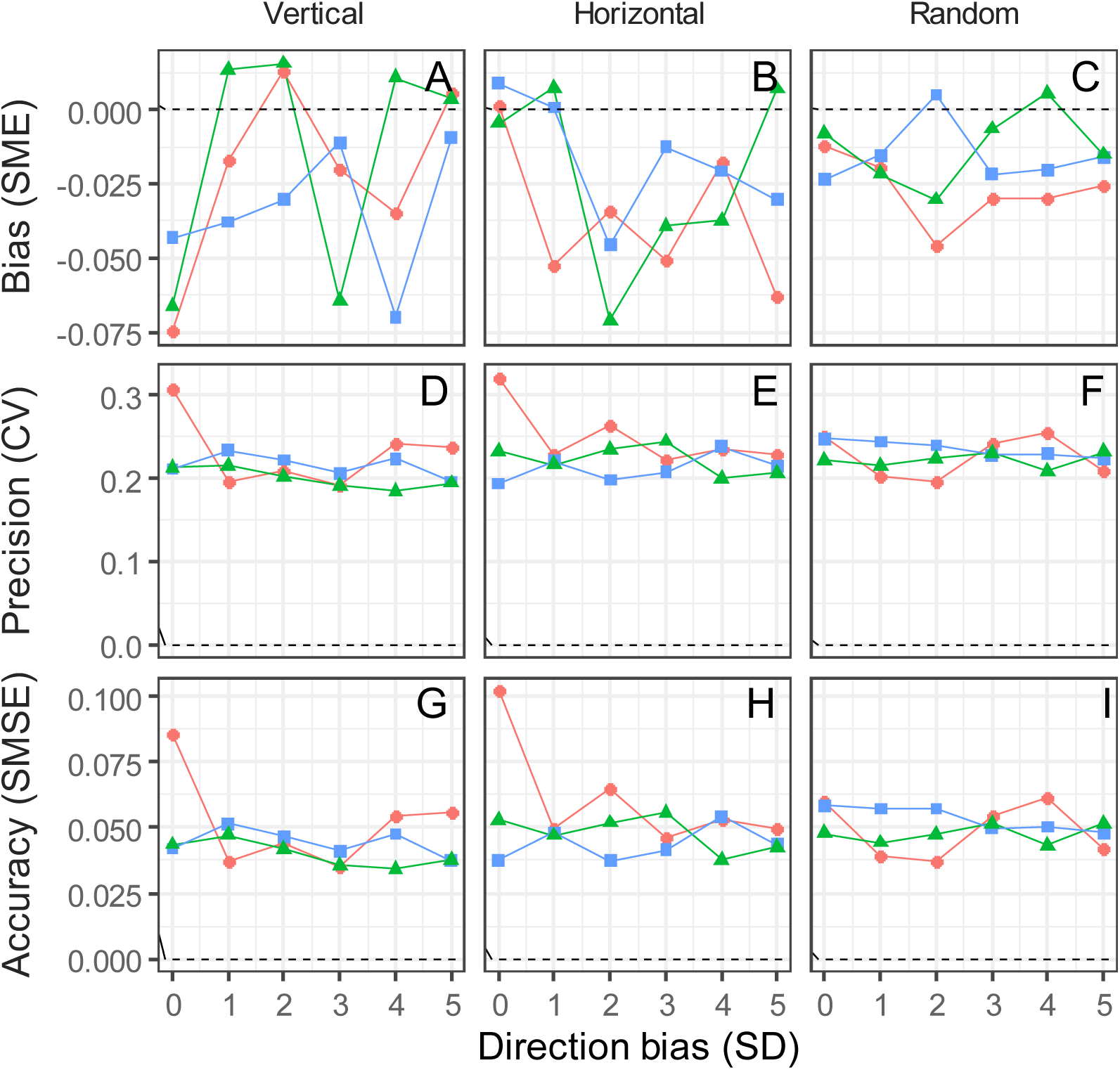
Effects of sampling strategies and animal movement bias on the performance of the 3D density estimation method. Columns represent the orientation of detectors: all vertical (left), all horizontal (middle) or all random (right). Symbols show the distribution of detectors: regular spacing on a 2D plane (circles), regular spacing in 3D throughout the sampling cube (triangles), and random distribution in 3D (squares). Parameters are as described in the text, using in particular a vertical direction component drawn at random from a truncated normal distribution centred at 0 with standard deviations between 0 and 5 to show direction bias.

## Discussion

We have outlined a method to use remote detectors such as underwater cameras or acoustic sensors to estimate population density from an encounter rate. The underlying random encounter model was originally proposed and tested as a density estimator for species moving in a two-dimensional terrestrial environment by Rowcliffe et al. (2008), and our calculations now allow its adaptation to species that move in three dimensions such as fishes, birds and bats. The basic requirements regarding detector specifications and information on movement speed remain the same as for the two-dimensional case.

Simulations show good performance of the estimator, low levels of bias, and a high degree of precision. Such consistent performance was expected when all assumptions were met, which may not be the case in real-life applications. We showed, however, that the method is robust to violations of assumptions. There was very little effect of increased variance in speed among individuals, or of bias in direction. Furthermore, our simulation results suggest that any effect could be limited or altogether eliminated simply by orienting detectors in different directions, even when detectors are placed on a single plane (on the ground, for example).

Performance is significantly influenced by sampling effort as it relates to the real density, in terms of both time and number of detectors. At low densities especially, insufficient effort could result in an over- or underestimation of density. An advantage of using a method based on movement models is that researchers can use the same framework to calculate the effort needed beforehand. Rearranging eq. 1 and substituting density and speed with prior information allows calculating an expected capture frequency. We saw no bias at higher densities, but we expect that in practice extremely high densities (e.g. in fish schools) may prevent properly counting individuals, resulting in underestimated densities. In these cases, researchers could first estimate a density of groups and obtain overall density by multiplying this estimate by an independently calculated mean group size (Rowcliffe et al. 2008).

Our method can be applied in cases where a lack of individual markings impedes the use of mark-recapture techniques. It is also an improvement over indices of relative abundance such as the maximum or mean number of conspecifics in a single frame (Schobernd, Bacheler, & Conn, 2014; Sherman et al., 2018). These metrics are commonly used to analyse footage from baited-remote-underwater-video-stations (BRUVS), but can underestimate true abundance (Cappo, Harvey, Malcolm, & Speare, 2003; Stobart et al., 2015). We know of no similar tools to estimate abundance from acoustic detectors. These sensors allow to identify species and count passages through the detection zone, so our method is also applicable with these technologies, provided the detection zone and mean species speed can be accurately measured.

For many species, mean speed will not be immediately available in the literature but could be approximated with additional measurements. For example, using two cameras in a stereo arrangement, the footage from both could be used to estimate speed (Somerton, Williams, & Campbell, 2017; Williams, Rooper, & Towler, 2010) and – using only one of the two cameras – population density in the same study.. Movement If detectors are deployed for several days the estimation of mean speed must include periods of inactivity (see Carbone, Cowlishaw, Isaac, & Rowcliffe, 2005). Ideally, surveys should be conducted at the same time of day when working in different sites, during the species’ daily activity peak.

Additional to speed, an exact characterization of the detection zone is required to estimate density accurately. This zone is determined first by the opening angle (for acoustic detectors), or the horizontal and vertical fields of vision (for cameras), usually given by the manufacturer. However, action cameras commonly used in remote underwater surveys have wide-angle lenses, which distort the image. Because of this, the diagonal field of vision will not correspond to the angle calculated assuming a rectilinear lens (see Appendix A3). The additional area in the projection due to the distortion should be small, so we suggest assuming a rectilinear lens for consistency.

The second element needed to characterize the detection zone is the detection radius. Unlike the opening angles the detection radius is influenced by environmental variables. For example, for underwater cameras, detection distance depends on visibility (i.e. turbidity), which should be considered when comparing densities across sites. Similarly, for acoustic detectors, atmospheric conditions like temperature and humidity affect how far an acoustic signal travels, effectively influencing the detection distance for birds and bats (Lawrence & Simmons, 1982; Snell-Rood, 2012). In both aquatic and aerial surveys, physical obstacles such as vegetation will also limit detectability; for instance, detection of bats with low-frequency calls (25 kHz) is significantly hindered by habitat structure (Patriquin, Hogberg, Chruszcz, Barclay, & Barclay, 2003; Weller & Zabel, 2002).

Environmental variables also interact with species-specific traits, generating different detection distances even under comparable environmental conditions. For example, cameras can detect larger species further than smaller species, and acoustic detectors will detect species with lower frequency or more intense calls at greater distances from than species with high-frequency calls (Jakobsen, Brinkløv, & Surlykke, 2013; Lawrence & Simmons, 1982; Snell-Rood, 2012; Surlykke & Kalko, 2008). Given the multiple factors that influence detection distance, we suggest an ad-hoc calculation for every system, for example placing a model of the study species progressively further from a camera until it is no longer recognizable. With acoustic detectors the same can be done using a speaker at a range of distances.

Our calculations are performed for individual detectors, but, as our simulations show, the best results are obtained when averaging across multiple detectors. Improved performance using more detectors is to be expected, as averaging across detectors minimizes possible sampling errors (Rowcliffe et al., 2008). Using more detectors also reduced variability across trials, implying that less effort is required to obtain accurate density estimates. Sampling designs should seek to maximize the number of encounters with a target species. using a number and spacing of detectors that capture the movement of the species of interest, while maintaining independence across detectors (Keiter et al., 2017).

Finally, we note that detection zones can also be affected by placement. Cameras placed in shallow streams, for example, could have their detection zone cut at the top by the surface and at the bottom by the substrate. In these cases, the resulting video frames can be cropped so that neither the ground nor the surface are visible, and the fields of vision and capture frequency recalculated accordingly (this would be equivalent to having a camera with narrower field of vision). If topography permits, one can prevent the issue of an incomplete detection zone by placing the camera in mid-water such that neither the water surface nor the bottom are visible. This would be a departure from current designs that set cameras on the ground but would avoid the issue of cropping the detection zone. For benthic species or very shallow streams none of these solutions might be feasible, and we would recommend using a 2D approach.

In summary, we have proposed a method for estimating density in three dimensions using data from remote detectors, which can be used in ecological and conservation research and as a monitoring tool. The description of detection zones provided will be useful in translating other density estimation methods that are also based on the ideal gas model (Campos-Candela et al., 2018; Moeller, Lukacs, & Horne, 2018). There is an extensive and growing literature characterizing sampling requirements for camera-traps in two dimensions, but such considerations are practically non-existent for species moving in three dimensions (Burton et al., 2015; O’Connell, Nichols, & Karanth, 2010). Our analyses allow extending these considerations to three dimensions, and it is our hope that our work will prompt new study designs and applications of remote detection methods to study a broader range of species and environments. We lay out the theoretical foundation of the method but recognize that it will require empirical validation. The sampling of aquatic, airborne, and arboreal species each comes with intrinsic challenges and field trials must be conducted to confirm that the method performs in real conditions as well as predicted by simulations. Given the existing camera-trapping literature and the use of underwater cameras to estimate abundance, we believe the application of our method will be more straightforward in underwater censuses. The application to acoustic detectors will require further work to characterize the detection zone, as currently there is no quantitative way to measure the detection distance under different environmental conditions. Thus, the requirements of our method open new avenues of research in remote detection.

## Acknowledgments

PKM is grateful for support from an NSERC (Natural Sciences and Engineering Research Council of Canada) Discovery Grant, CFI (Canada Foundation for Innovation) John R. Evans Leader Funds, and MRIS Ontario Research Funds. Funding by FRQNT (Fonds de recherche nature et technologie) to RAC and an NSERC Discovery Grant to NEM is gratefully acknowledged.

## Authors’ contributions

RAC, PKM and JSVS conceived the original concept. JSVS and PKM derived the equations. JSVS conducted simulations and led the writing of the manuscript. All authors contributed critically to the drafts and gave final approval for publication.

## Data accessibility

The code for the simulations and to reproduce figures is available on github in the following repository: juansvs/3D_DensityEstimation

