## Supplementary Material for "Estimating animal density in three dimensions using capture-frequency data from remote detectors"

**Appendices**

A1. Calculation of the height of a spherical cap

A2. Surface area of a spherical rectangle

A3. Calculation of fields of vision and lateral surface angles of cropped detectors (cameras)

A4. Non-uniformly distributed approach directions: derivation of equations 10 and 11

A5. Step-by-step guide to using the density estimation formula

**A1. Calculation of the height of a spherical cap**

The top section of an acoustic – i.e. conical – detection zone is a spherical cap. The surface area of this cap is determined by the radius of the corresponding sphere (here, given by the cone's slant height *s*) and the cap's height *h*. The following trigonometric calculations yield the formula for *h* that is used in equation 5 of the main text (cf. Figure S1 for notation):

$$AC = s\cos\frac{\phi}{2} \text{and } AB = AD = s$$

As such,

$$h = DC = AD - AC = s\left(1 - \cos\frac{\phi}{2}\right) \qquad (S1.1)$$

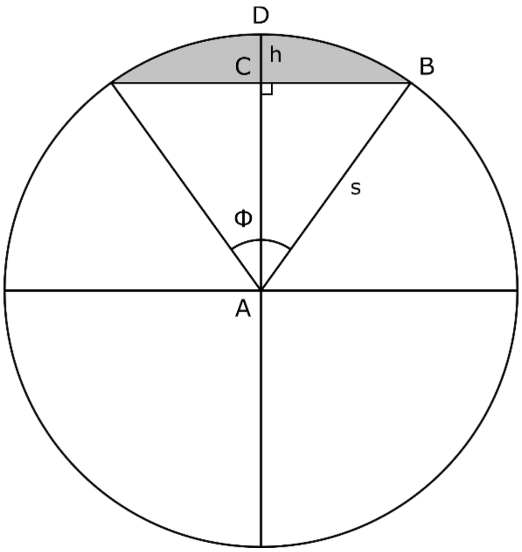


**Figure S1. Cross section of an acoustic detection zone, showing the measurements needed to calculate the height *h* of the spherical cap (shaded region). The detector is located at the vertex A, which is also the center of the sphere of radius *s* that the spherical cap is a part of. The angle $\phi$ is the opening angle of the detector. See text for calculations.**

**A2. Surface area of a spherical rectangle**

The top section of a camera-trap's detection zone is a spherical rectangle whose surface area can be calculated by considering the proportion of the surface area of the sphere of radius *s* that it covers. To determine this proportion, we use the concept of solid angles. Intuitively, a solid angle can be understood as a measure for the amount of the field of view that is covered by a given object from some particular observation point. Formally, an object's solid angle is defined as the area of the unit sphere that is blocked by the object from the view of an observer located at the center of the sphere. In our case, the object is the rectangular frame that is determined by the horizontal and vertical fields of vision of the camera, $\kappa$ and $\lambda$ (Fig. S2), which has solid angle Ω, calculated as follows (see proof below):

$$\Omega = 4 \arcsin \frac{\tan\frac{\kappa}{2} \tan\frac{\lambda}{2}}{\sqrt{\tan^2\frac{\kappa}{2} + 1}\sqrt{\tan^2\frac{\lambda}{2} + 1}} \qquad (S2.1)$$

The area of the spherical rectangle atop a camera's detection zone is then:

$$S_R = 4\pi s^2 \frac{\Omega}{4\pi} = s^2\Omega \qquad (S2.2)$$

because solid angles are measured in steradians (sr), a unitless quantity, and the maximum solid angle for the complete unit sphere is 4π sr. The proportion of the unit sphere occupied by the solid angle of the camera frame is then $\frac{\Omega}{4\pi}$. This proportion is multiplied by the surface area of the sphere of radius *s*, which equals $4\pi s^2$, to obtain the surface area of the spherical rectangle as in equation S2.2.

To see the validity of equation S2.2, we note that the solid angle of a rectangle with side lengths *a* and *b* that is located at a distance *c* in front of an observer (Fig. S2) is given by

$$\Omega = 4 \arcsin \frac{ab}{\sqrt{a^2 + c^2}\sqrt{b^2 + c^2}} \qquad (S2.3)$$

Khadjavi (1968), and that:

$$a = c \tan\frac{\kappa}{2}; \text{ and } \quad b = c \tan\frac{\lambda}{2} \tag{S2.4}$$

where *κ* and *λ* are the horizontal and vertical fields of vision (Fig. S2). Substituting *a* and *b* from equation (S2.4) into (S2.3) we obtain equation (S2.1).

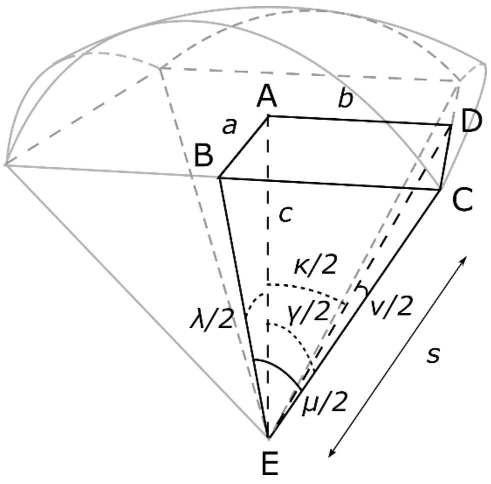


**Figure S2: Section of the cropped detection zone of camera traps. The dark sections are used in the calculation of the vertical and diagonal fields of vision (FOVs) *λ* and *γ*, and the lateral angles *μ* and *ν*. The rectangle ABCD, with side lengths *a* and *b*, is one quarter of the image frame delimited by the vertical and horizontal FOVs, and is located a distance *c* from the detector at E.**

### A3. Calculation of fields of vision and lateral surface angles of cropped detectors (cameras)

Application of our methods requires knowing the horizontal, vertical, and diagonal fields of vision of the camera ($\kappa$, $\lambda$ and $\gamma$), as these are all required to calculate the opening angles ($\mu$ and $\nu$) of the disk sectors that make up the sides of the camera detection zone. However, camera manufacturers often only provide the horizontal field of vision (FOV). Here we show how to calculate all the unknown angles based on a single known angle and the aspect ratio of the image frame. First, in Appendix A3a, we show how to determine the vertical FOV from the horizontal FOV if it is not already provided by the manufacturer; in Appendix A3b, we show how to determine the diagonal FOV from the horizontal and vertical FOVs; and in Appendix A3c, we show how to determine the lateral opening angles using all the FOVs.

#### *A3a. Calculation of the vertical FOV from the horizontal FOV*

Some manufacturers provide both the horizontal and vertical FOVs, $\kappa$ and $\lambda$. If only the horizontal FOV is provided, the vertical FOV can be obtained from the horizontal FOV and the aspect ratio, $q$, as:

$$\tan\frac{\lambda}{2} = q\tan\frac{\kappa}{2} \qquad (S3.1)$$

To see this, consider the pyramid ABCDE in figure S2, which represents one quarter of the base of the cropped detection zone. Defining $AB = b$, $AD = a$, and $AE = c$, we obtain $a = c\tan\frac{\lambda}{2}$ and $b = c\tan\frac{\kappa}{2}$, given that $\widehat{AED} = \frac{\kappa}{2}$ and $\widehat{AEB} = \frac{\lambda}{2}$. Because we also have $b = qa$, where $q$ is the user-selected aspect ratio (most commonly 3/4 or 9/16), we obtain (S3.1).

#### *A3b. Calculation of the diagonal FOV from the horizontal and vertical FOVs*

If a diagonal field of vision of a camera detector is not provided by the manufacturer, it can be determined from the detector's horizontal and vertical fields of vision, $\kappa$ and $\lambda$. Even if the diagonal FOV is provided, we suggest using the calculations here for consistency with the calculations of other angles. For a rectilinear, non-distorted lens, $\gamma$ is given by:

$$\gamma = 2\arctan\sqrt{\tan^2\frac{\lambda}{2} + \tan^2\frac{\kappa}{2}} \tag{S3.2}$$

To see this, consider the triangle ABC in Fig. S2, and note that $AB^2 + BC^2 = AC^2$. Furthermore, we have $AB = c\tan\frac{\lambda}{2}$, $BC = c\tan\frac{\kappa}{2}$, and $AC = c\tan\frac{\gamma}{2}$, from which it follows that

$$\left(c\tan\frac{\lambda}{2}\right)^2 + \left(c\tan\frac{\kappa}{2}\right)^2 = \left(c\tan\frac{\gamma}{2}\right)^2$$

$$\Rightarrow \sqrt{c^2\left(\tan^2\frac{\lambda}{2} + \tan^2\frac{\kappa}{2}\right)} = c\tan\frac{\gamma}{2}$$

ctan2λ2+tan2κ2=ctanγ2#(Sand

and thus, equation (S3.2).

*A3c. Calculation of the lateral opening angles μ and v from the fields of vision*

The sides of the base of the cropped detection zone are disk sectors, whose surface areas are calculated by multiplying the total surface of a disk of radius *s* by the proportion of the disk that is occupied by the disk sectors. These proportions are given by $\frac{\mu}{2\pi}$ and $\frac{\nu}{2\pi}$, respectively, where $\mu$ and $\nu$ are the frontal and lateral opening angles of the detection zone (Fig. S2), and which can be calculated from the diagonal, vertical, and horizontal fields of vision, $\gamma$, $\lambda$, and $\kappa$ as:

$$\nu = 2\arccos\frac{\cos\frac{\gamma}{2}}{\cos\frac{\kappa}{2}} \text{ and } \mu = 2\arccos\frac{\cos\frac{\gamma}{2}}{\cos\frac{\lambda}{2}} \tag{S3.4}$$

To see this, note that $\frac{\mu}{2} = \widehat{DEC}$ and $\frac{\nu}{2} = \widehat{BEC}$ in Figure S3. DEC is a right triangle so $DE = CE \cos\frac{\nu}{2}$. Moreover, $AE = DE \cos\frac{\kappa}{2}$ from which it follows that

$$AE = CE \cos\frac{\nu}{2}\cos\frac{\kappa}{2} \qquad (S3.5)$$

We also have EAC is a right triangle, so:

$$AE = CE \cos\frac{\gamma}{2} \qquad (S3.6)$$

Equating (S3.5) and (S3.6) we obtain:

$$CE \cos\frac{\nu}{2}\cos\frac{\kappa}{2} = CE \cos\frac{\gamma}{2}$$

$$\cos\frac{\nu}{2} = \frac{\cos\frac{\gamma}{2}}{\cos\frac{\kappa}{2}}$$

and thus, the first equation S3.4. The second equation is obtained in the same manner, only using $\lambda$ instead of $\kappa$.

**A4. Non-uniformly distributed approach directions: derivation of equation 12.**

In natural systems it is likely there will be some bias in movement, which invalidates the assumption of equally likely directions of approach to the detection zone. Our simulations show that any error in the estimation of density caused by such biases can be prevented by setting up multiple detectors facing in different directions. However, if only a few detectors are available or if the sampling region is not large enough to set multiple detectors independently, strong direction bias should be addressed in the calculation of the mean profile area.

*A4a. Derivation of formula in equation 12*

To calculate the mean projected area of a detection zone, $\hat{p}$, for the case of non-uniformly distributed approach directions, we derive formulae for the detection zone's projected area, $p(\omega, \theta)$, for all possible directions of approach $(\omega, \theta)$, and then weight these according to the probability distribution of approach directions $P(\omega, \theta)$, which equals $P(\omega)P(\theta)$ in case of independence, as follows:

$$\hat{p} = \iint P(\omega)\, P(\theta)\, p(\omega, \theta)\, d\theta\, d\omega \qquad (S4.1)$$

As seen in Fig. 3, if the detector is oriented along the *z* axis, $\theta$ is the azimuth angle on the (x, y) plane, between 0 and 2π. The angle $\omega$ is the angle with respect to the *z* axis, from 0 to π. Given the radial symmetry of acoustic detectors, the profile of the detection zone varies only with respect to ω, so we obtain:

$$\hat{p} = \int_0^{\pi} P(\omega)\, p(\omega)\, d\omega \qquad (S4.2)$$

The profile $p(\omega)$ will have a different shape depending on the approach direction and which parts of the detection zone are visible to an animal given its approach direction. Thus, the profiles can be visualized as combinations of three components: an ellipse, a "hat," and an arc (areas I, II and III in

Fig. 3), corresponding to the projections of the base of the conic section, the sides of the cone, and the spherical cap, respectively. There are four possible combinations of these areas:

$$p(\omega) = \begin{cases} a_e(\omega), & \sin\omega \le \sin\frac{\phi}{2} \text{ and } \sin\omega \le \cos\frac{\phi}{2} \\ a_e(\omega) + a_d(\omega), & \sin\omega \le \sin\frac{\phi}{2} \text{ and } \sin\omega > \cos\frac{\phi}{2} \\ a_e(\omega) + a_h(\omega), & \sin\omega > \sin\frac{\phi}{2} \text{ and } \sin\omega \le \cos\frac{\phi}{2} \\ a_e(\omega) + a_d(\omega) + a_h(\omega), & \sin\omega > \sin\frac{\phi}{2} \text{ and } \sin\omega > \cos\frac{\phi}{2} \end{cases} \qquad (S4.3)$$

The first case applies when only the ellipse is visible, which occurs when the animal is approaching the detector almost directly from the front or the back. The second and third cases apply when the ellipse is visible, along with either the spherical cap (second case) or the conic base (third case). The fourth case is applicable when all three parts are visible, which occurs when animals approach the detection zone from the sides. Only three of these four cases will be applicable for a given detector, and this depends on its opening angle: for $\phi > \frac{\pi}{2}$ there is no direction of approach where the cone is visible, and the dome is not. For $\phi < \frac{\pi}{2}$ there is no direction where the dome is visible, and the cone is not (Fig. 3).

The area $a_e$ can be calculated directly with the equation for the area of an ellipse (see equation S4.4). The other two areas, $a_h$ and $a_d$, are calculated as a difference of integrals. First, we determine the equations that define the boundaries of the arc, the hat and the ellipse. Then, we find the intersections between the curves. Finally, we integrate the difference between the equations of the arc or the hat and the ellipse, and the intersections. This procedure follows Pennell and Deignan (1989) who studied the projected area of a cone.

The result for a cone was already given by Pennell and Deignan (1989) in terms of the height of the cone *c* and the radius at the base *r*. Noting that $c = s\cos\frac{\phi}{2}$ and $r = s\sin\frac{\phi}{2}$, their result can be expressed as:

$$p(\omega) = \begin{cases} \pi\left(s\sin\frac{\phi}{2}\right)^2 |\cos\omega|, & |\tan\omega| \leq \tan\frac{\phi}{2} \\ \pi\left(s\sin\frac{\phi}{2}\right)^2 |\cos\omega| + a_h(\omega), & |\tan\omega| > \tan\frac{\phi}{2} \end{cases} \quad (S4.4)$$

The difference of integrals between 0 and *x** is one half of the total area, we multiply this difference by two to calculate the total area of the hat , where *x** is the abscissa of the intersection between the curves (Pennell and Deignan 1989). We have $x^* = \tan\frac{\phi}{2}\sqrt{\left(s\cos\frac{\phi}{2}\right)^2 - \left(s\sin\frac{\phi}{2}\right)^2\cot^2\omega}$, and $a_h$ is given by:

$$a_h(\omega) = \left(2s\cos\frac{\phi}{2}\sin\omega\right)x^* - \frac{(x^*)^3\left(s\cos\frac{\phi}{2}\sin\omega\right)}{\left(s\sin\frac{\phi}{2}\right)^2}$$
$$-|\cos\omega|\left[\left(s\sin\frac{\phi}{2}\right)^2\sin^{-1}\left(\frac{x^*}{s\sin\frac{\phi}{2}}\right) + x^*\sqrt{\left(s\sin\frac{\phi}{2}\right)^2 - (x^*)^2}\right] \quad (S4.5)$$

Here, we follow the same steps to expand equation S4.4 to include the area of the projection of the spherical cap (area II in Fig 3). The corresponding arc can be described using the equation for a circle of radius *s*, centered at the projection of the vertex of the cone $\left(0, s\cos\frac{\phi}{2}\sin\omega\right)$, as follows:

$$(x-0)^2 + \left(y - s\cos\frac{\phi}{2}\sin\omega\right)^2 = s^2$$

$$\Rightarrow \left(y - s\cos\frac{\phi}{2}\sin\omega\right)^2 = s^2 - x^2$$

$$\Rightarrow y - s\cos\frac{\phi}{2}\sin\omega = \pm\sqrt{s^2 - x^2}$$

$$\Rightarrow y = s\cos\frac{\phi}{2}\sin\omega \pm \sqrt{s^2 - x^2} \quad (S4.6)$$

Here we only use the equation for the lower arc. At the point of intersection between the arc and the ellipse, their curves are tangent, so we can find the coordinates by equating their slopes. The slope for the arc is $y' = \frac{x}{\sqrt{s^2-x^2}}$. From Pennell and Deignan (1989), we further know that the slope of the bottom half of the ellipse is $y' = \frac{x|\cos\omega|}{\sqrt{s^2\sin^2\frac{\phi}{2}-x^2}}$. We therefore have:

$$\frac{x|\cos\omega|}{\sqrt{s^2\sin^2\frac{\phi}{2}-x^2}} = \frac{x}{\sqrt{s^2-x^2}}$$

$$\Rightarrow |\cos\omega| = \frac{\sqrt{s^2\sin^2\frac{\phi}{2}-x^2}}{\sqrt{s^2-x^2}}$$

$$\Rightarrow \cos^2\omega = \frac{s^2\sin^2\frac{\phi}{2}-x^2}{s^2-x^2}$$

$$\Rightarrow \cos^2\omega\,(s^2-x^2) = s^2\sin^2\frac{\phi}{2}-x^2$$

$$\Rightarrow s^2\cos^2\omega - x^2\cos^2\omega + x^2 = s^2\sin^2\frac{\phi}{2}$$

$$\Rightarrow x^2(1-\cos^2\omega) = s^2\left(\sin^2\frac{\phi}{2}-\cos^2\omega\right)$$

$$\Rightarrow x^2 = \frac{s^2\left(\sin^2\frac{\phi}{2}-\cos^2\omega\right)}{1-\cos^2\omega}$$

$$\Rightarrow x = \pm\sqrt{\frac{s^2\left(\sin^2\frac{\phi}{2}-\cos^2\omega\right)}{1-\cos^2\omega}}$$

$$\Rightarrow x^* = \pm\, s\sqrt{\frac{\sin^2\frac{\phi}{2}-\cos^2\omega}{1-\cos^2\omega}} \qquad (S4.7)$$

We now integrate the difference between the arc and the ellipse (given by $y = -|\cos\omega|\sqrt{s^2\sin^2\frac{\phi}{2}-x^2}$) from 0 to $x^*$. The full area $a_e$ is two times the integral:

$$a_d(\omega) = 2\int_0^{x^*} s\cos\frac{\phi}{2}\sin\omega - \sqrt{s^2 - x^2} - \left(-|\cos\omega|\sqrt{s^2\sin^2\frac{\phi}{2} - x^2}\right)dx$$

$$\Rightarrow a_d(\omega) = 2\int_0^{x^*} s\cos\frac{\phi}{2}\sin\omega - \sqrt{s^2 - x^2} + |\cos\omega|\sqrt{s^2\sin^2\frac{\phi}{2} - x^2}\,dx$$

$$\Rightarrow a_d(\omega) = 2\left(s\cos\frac{\phi}{2}\sin\omega\ x^* - \int_0^{x^*}\sqrt{s^2 - x^2}\,dx + |\cos\omega|\int_0^{x^*}\sqrt{s^2\sin^2\frac{\phi}{2} - x^2}\,dx\right)$$

$$\Rightarrow a_d(\omega) = 2\left(s\cos\frac{\phi}{2}\sin\omega\ x^* - \frac{1}{2}\left(x^*\sqrt{s^2 - x^{*2}} + s^2\arctan\frac{x^*}{\sqrt{s^2 - x^{*2}}}\right)\right.$$

$$\left. + |\cos\omega|\frac{1}{2}\left(x^*\sqrt{s^2\sin^2\frac{\phi}{2} - x^{*2}} + s^2\sin^2\frac{\phi}{2}\arctan\frac{x^*}{\sqrt{s^2\sin^2\frac{\phi}{2} - x^{*2}}}\right)\right)$$

*A4b. Approach angles where each formula in equation 12 applies*

We will now focus on determining the cases where each component is visible. Pennell and Deignan (1989) gave the cases where area I, the ellipse, and area III, the "hat" are visible (see Equation S15). As ω increases, the arc becomes visible when the direction of approach is perpendicular to the near side of the conic section (direction OP in Fig. S3). We know $\widehat{OQP} = \frac{\phi}{2}$, thus

$$\widehat{QOP} = \pi - \left(\frac{\pi}{2} + \frac{\phi}{2}\right)$$

$$\widehat{QOP} = \frac{\pi}{2} - \frac{\phi}{2}$$

The projection of the spherical cap disappears from view when the direction of approach is perpendicular to the far side of the cone; this is, when the direction is parallel to OR (Fig. S4). The triangle ORQ is similar to OPQ, so $\widehat{QOP} = \widehat{QOR} = \frac{\pi}{2} - \frac{\phi}{2}$. The opposite angle is also the difference between π and the angle ω, we can therefore calculate ω as follows:

$$\pi - \omega = \left(\frac{\pi}{2} - \frac{\phi}{2}\right)$$

$$\omega = \pi - \left(\frac{\pi}{2} - \frac{\phi}{2}\right)$$

$$\omega = \frac{\pi}{2} + \frac{\phi}{2}$$

The arc is therefore evident in the projection if:

$$\frac{\pi}{2} - \frac{\phi}{2} < \omega < \frac{\pi}{2} + \frac{\phi}{2}$$

$$\Rightarrow \sin\omega > \sin\left(\frac{\pi}{2} - \frac{\phi}{2}\right)$$

$$\Rightarrow \sin\omega > \cos\frac{\phi}{2}$$

Thus, we have shown the conditions for equation (S4.4):

$$p(\omega) = \begin{cases} a_e(\omega), & \sin\omega \le \sin\frac{\phi}{2} \text{ and } \sin\omega \le \cos\frac{\phi}{2} \\ a_e(\omega) + a_d(\omega), & \sin\omega \le \sin\frac{\phi}{2} \text{ and } \sin\omega > \cos\frac{\phi}{2} \\ a_e(\omega) + a_h(\omega), & \sin\omega > \sin\frac{\phi}{2} \text{ and } \sin\omega \le \cos\frac{\phi}{2} \\ a_e(\omega) + a_d(\omega) + a_h(\omega), & \sin\omega > \sin\frac{\phi}{2} \text{ and } \sin\omega > \cos\frac{\phi}{2} \end{cases}$$

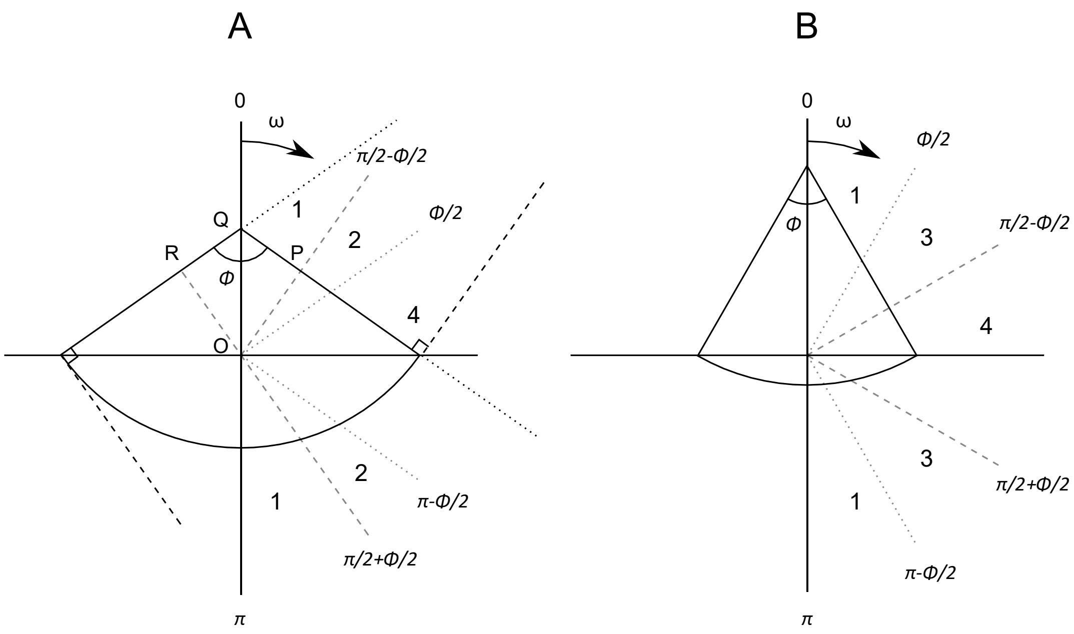


**Figure S3. Longitudinal cross section of a conical detection zone, depicting the angles of approach ω that define the different cases for calculating the projected area. The dotted lines are parallel to the sides and therefore their direction is tangent to the sides. As such, they mark the limit angles between which the conical hat ($a_h$) is visible in the projection. The dashed lines are perpendicular to the sides of the cone; their direction is tangent to the spherical cap and as such they mark the angles between which the arc ($a_d$) is visible. Therefore, in 1, only the ellipse is visible; in 2 the ellipse and the arc are visible; in 3 the ellipse and the hat are visible; in 4 all three areas are visible. The gray lines show the same directions, translated to the origin to see the relationship between the angles. The two figures show the case of a detection zone with opening angle ϕ greater (A) or less than π/2 (B). The points O, P, and Q are used in the calculations of *ω* for the different cases.**

#### *A4c. Determination of frequency distribution of angles of approach with cameras*

To calculate the weighted mean profile area, we require the frequency distribution of angles of approach to the detection zone. Our coordinate reference system is defined by the detector, and the direction of an individual can be characterized by two angles: $\theta$, the azimuth angle perpendicular to the direction of the detector, and $\omega$, the angle with respect to the direction of the detector (Fig. S4). Two cameras can be set up overlooking the same focus area, but perpendicular to each other to determine these two angles. Over time with sufficient individuals coming into the detection zone a frequency distribution can be approximated. Alternatively, one can characterize the direction of an animal in three dimensions using a stereo arrangement (two cameras side by side looking in slightly different directions, e.g. Somerton, Williams, & Campbell, 2017), and this direction can then be decomposed into the two angles $\omega$ and $\theta$.

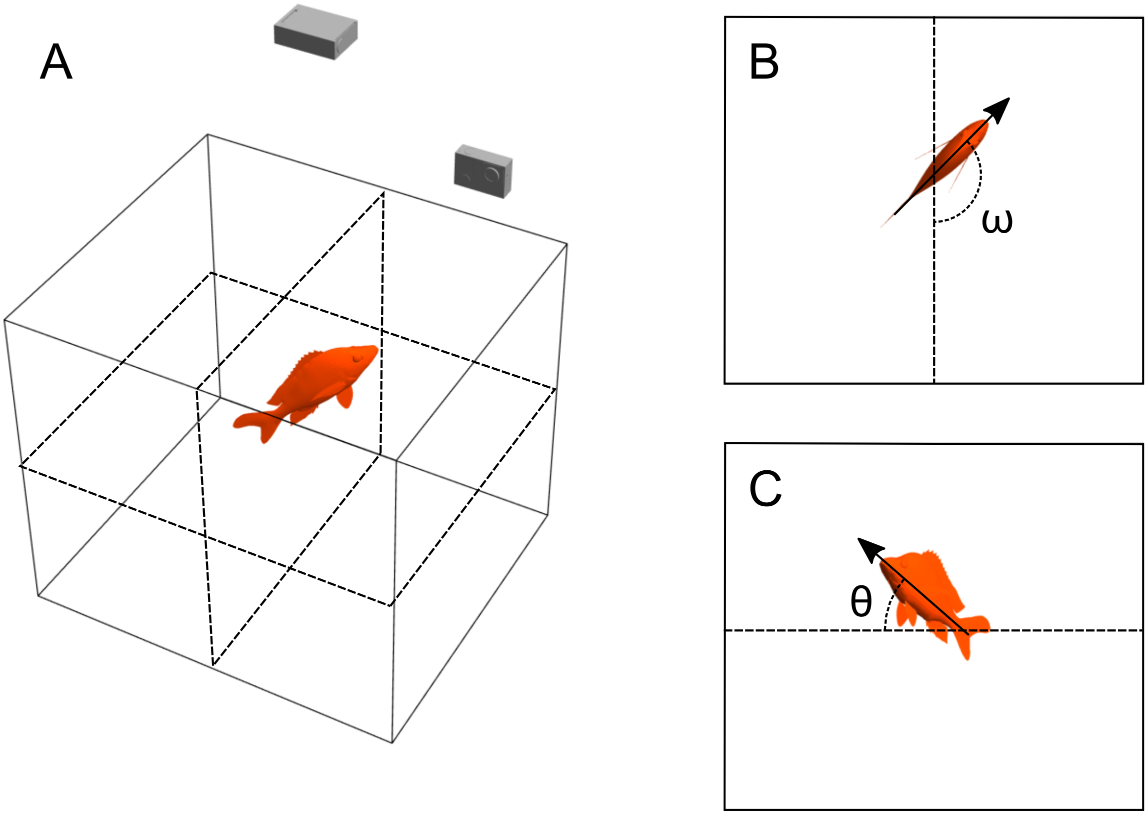


**Figure S4. Two-camera arrangement for determining the frequency distribution of approach angles (A). Two cameras perpendicular to each other allow decomposing the direction of an individual into the two angles $\theta$ and $\omega$ necessary to characterize its three-dimensional direction. The arrows show the direction the animal is travelling, and the dashed lines are reference lines with respect to the camera on the side. The insets show the image captured by the cameras from the top (B), where the dashed line represents the line of sight of the other**

**camera, and the view of the camera on the side (C), where the dashed line is a horizontal reference for the angles.**

### A5. Step-by-step guide to using the density estimation formula

Here we show the steps necessary to estimate density with the 3D REM method with a numerical example. First, we use the detector's specifications to define its detection zone and calculate its profile area. Second, we deploy the detector, and obtain a detection frequency. The last step is to calculate the density using these two previously calculated values.

#### *A5a. Calculation of the profile area*

We will assume a single camera, for which we know the horizontal FOV ($\kappa$) is 120° (2π/3 rad) but we don't know the vertical or diagonal FOV. We must then use the formulas in appendix A3 to calculate the missing angles. Say we set the camera to have an aspect ratio $q$ of 3/4. We can then use eq. S3.1 to calculate the vertical FOV $\lambda$:

$$\tan\frac{\lambda}{2} = q\tan\frac{\kappa}{2}$$

$$\tan\frac{\lambda}{2} = \frac{3}{4}\tan\left(\frac{1}{2}\times\frac{2\pi}{3}\right)$$

$$\frac{\lambda}{2} = \arctan 1.30$$

$$\lambda = 1.83\ rad = 104.85°$$

Once we have both the horizontal and vertical FOV we can use them to calculate the diagonal FOV, using eq. S3.2:

$$\gamma = 2\arctan\sqrt{\tan^2\frac{\lambda}{2} + \tan^2\frac{\kappa}{2}}$$

$$\gamma = 2\arctan\sqrt{\tan^2\frac{1.83}{2} + \tan^2\frac{\pi}{3}}$$

$$\gamma = 2.28\,rad = 130.63°$$

After obtaining all the FOV angles, we can calculate the lateral opening angles $\mu$ and $\nu$, using eq. S3.4:

$$\nu = 2\arccos\left(\frac{\cos\frac{\gamma}{2}}{\cos\frac{\kappa}{2}}\right)$$

$$\nu = 2\arccos\left(\frac{cos\frac{2.28}{2}}{cos\frac{\pi}{3}}\right)$$

$$\nu = 1.16\,rad = 66.46°$$

By the same method we obtain $\mu = 1.63\,rad = 93.39°$. These two angles are then used to calculate the lateral surface area of the detection zone as a function of the detection distance, using eq. 7:

$$S_L = 2S_\nu + 2S_\mu = s^2(\mu + \nu)$$

$$S_L = s^2(1.63 + 1.16) = 2.79\,s^2$$

The other section of the detection zone is the spherical rectangle. We can calculate its solid angle in steradians (sr) using eq. S2.1:

$$\Omega = 4\arcsin\frac{\tan\frac{\kappa}{2}\tan\frac{\lambda}{2}}{\sqrt{\tan^2\frac{\kappa}{2}+1}\sqrt{\tan^2\frac{\lambda}{2}+1}}$$

$$\Omega = 4\arcsin\frac{\tan\frac{\pi}{3}\tan\frac{1.83}{2}}{\sqrt{\tan^2\frac{\pi}{3}+1}\sqrt{\tan^2\frac{1.83}{2}+1}}$$

$$\Omega = 3.34\,sr$$

The surface area of the section is then a function of $s$, obtained using eq. 8:

$$S_R = s^2\Omega$$

$$S_R = 3.34\ s^2$$

The total surface area is obtained by adding together the two surface areas calculated previously, and the mean profile area is one quarter of the sum (eq. 10):

$$\widehat{p_{cam}} = \frac{1}{4}(S_L + S_R)$$

$$\widehat{p_{cam}} = \frac{1}{4}(2.79s^2 + 3.34s^2)$$

$$\widehat{p_{cam}} = 1.53\ s^2$$

The profile area is proportional to the square of the detection distance *s*, which needs to be determined in the field, given the influence of environmental factors on visibility. Assuming a detection distance of 6 m, the mean profile area is $\widehat{p_{cam}} = 1.53 \times 6^2 = 55.08\ m^2$.

The procedure is much simpler for conical detectors. If we use the same angle of the diagonal FOV as the opening angle of the cone, the lateral area is given by eq.4:

$$S_C\ = \pi s^2 \sin\frac{\phi}{2}$$

$$S_C = 6^2\pi \sin\frac{2.28}{2}$$

$$S_C = 102.76\ m^2$$

And the surface area of the spherical cap is calculated with eq. 5:

$$S_S = 2\pi s^2 \left(1 - \cos\frac{\phi}{2}\right)$$

$$S_S = 2\pi \times 6^2 \left(1 - \cos\frac{2.28}{2}\right)$$

$$S_S = 131.74\ m^2$$

Adding $S_S$ and $S_C$ and dividing by four yields the mean profile area for a conical detection zone with similar characteristics as the camera (eq. 6):

$$\widehat{p_{aco}} = \frac{1}{4}(S_C + S_S)$$

$$\widehat{p_{aco}} = \frac{1}{4}(102.76 + 131.74)$$

$$\widehat{p_{aco}} = 58.63\ m^2$$

*A5b. Detection frequency and estimation of density*

We assume that the camera is deployed so that the detection zone is not limited, that is, far enough from the ground and the water surface. Say the camera was working for two hours, and during that time it captured a fish species 3 times. We have then a detection frequency $f = \frac{3}{2} h^{-1}$. We know beforehand that the mean speed of that study species is 1 m $s^{-1}$, or 3600 m $h^{-1}$. Given this information, and using the mean profile area of the camera's detection zone, we can calculate the three-dimensional density for that species using eq. 1:

$$D = \frac{f}{\hat{p}v}$$

$$D = \frac{1.5\ h^{-1}}{3600\ m.h^{-1}\ 55.08\ m^2}$$

$$D = 7.56 \times 10^{-6}\ m^{-3}$$

This is merely an illustration of the calculations necessary to estimate density, and the resulting value is not an accurate representation of any particular species.
